# Blood circulation in the tunicate *Corella inflata* (Corellidae)

**DOI:** 10.1101/029322

**Authors:** Michael W. Konrad

## Abstract

The ascidian tunicate *Corella inflata* is relatively transparent compared to other solitary tunicates and the circulatory system can be visualized by injecting high molecular weight fluorescein labeled dextran into the beating heart or the large vessels at the ends of the heart. In addition, after staining with neutral red the movement of blood cells can be followed to further define and characterize the circulatory system. The heart is a gently curved tube with a constriction in the middle and extends across the width of the animal. As in other tunicates, pumping is peristaltic and periodically reverses direction. During the abvisceral directional phase blood leaves the anterior end of the heart in two asymmetric vessels that connect to the two sides of the branchial basket (or pharynx), in contrast to the direct connection between the heart and the endostyle seen in the commonly studied tunicate *Ciona intestinalis.* In *Corella inflata* blood then flows in both transverse directions through a complex system of ducts in the branchial basket into large ventral and dorsal vessels and then to the visceral organs in the posterior of the animal. During the advisceral phase blood leaves the posterior end of the heart in vessels that repeatedly bifurcate to fan into the stomach and gonads. Blood speed, determined by following individual cells, is high and pulsatory near the heart, but decreases and becomes more constant in peripheral regions. Estimated blood flow volume during one directional phase is greater than the total volume of the animal. Circulating blood cells are confined to vessels or ducts in the visible parts of the animal and retention of high molecular weight dextran in the vessels is comparable to that seen in vertebrates. These flow patterns are consistent with a closed circulatory network.

## Introduction

Tunicates are the closest living relatives of vertebrates (Delsuc et al. 2006), and thus their study can play an important role in understanding vertebrate evolution. The tunicate *Ciona intestinalis* has many characteristics that have made it a model organism for study, and many hundreds of publications have described this species (Stolfi and Christiaen 2012). In particular, the development of the heart (Christiaen et al. 2009; Davidson 2007; Davidson et al. 2009; Stolfi et al. 2010), and by association the circulatory system, has been the subject of recent reports. However, the tunicate *Corella inflata* (Lambert et al. 1981) is more transparent than *C. intestinalis*, and this enables study of the circulatory system without the need of surgery to remove the mantle. In addition, in contrast to *C. intestinalis,* the body of *C. inflata* is rigid, and does not contract significantly when fluid is injected into vessels. In this report the circulatory system is defined by the path of blood cells or the space occupied by high molecular weight fluorescent tracer injected into the circulation.

## Methods

### Animal collection and care

*Corella inflata* Huntsman 1912 (Lambert et al. 1981) were collected in a marina in Sausalito, CA, USA (San Francisco Bay) from the side of a floating dock at a depth of 0.1 – 0.3 m. The present study is the result of observations on 94 animals over a period of 15 months.

Tunicates were kept in a seawater aquarium at temperatures of 16-20C with aeration, and used within 72 hours of collection. During injections and observations animals were submerged in seawater and pinned to a 5 mm thick gel of silicon (Slygard 184, Global-Industrial Corp) in a 7 x 7 cm acrylic box. Typically the top of the animal was only 1-3 mm below the surface of the seawater.

### Staining with neutral red

Tunicates were stained by immersion in 50 mL of a 0.1% solution of neutral red dye (Cynmar Corp.) in seawater for 10 minutes. They were then washed twice in 50 mL of seawater.

### Injection of fluorescent dextran

Fluorescent dextran (FITC labeled, 150,000 Daltons, Polysciences, Inc.) at a concentration of 5 mg/mL in seawater was injected using a 33-gauge x ½ inch hypodermic needle (TSK STERiJECT PRE-33013) connected by 8 cm of polyethylene tubing (PE/8, 1.2 mm ID, Scientific Commodities Inc.) to a 1 mL plastic tuberculin syringe (McKesson, 102-ST1C) with a total plunger travel of 57 mm. The syringe plunger was directly connected to a screw attached to a microprocessor controlled stepping motor with 200 steps per revolution. Rotation of the screw thus rotated the plunger to overcome static friction that could affect fluid delivery. One revolution advanced the screw 0.85 mm corresponding to 15 uL. Injections were controlled by joystick, with stepper motion recorded to a computer file. Typically 5 to 20 uL were injected over a 3 to 15 second period.

### Imaging of fluorescent dextran

Excitation was accomplished using a single LED (Luxeon V Star Blue, 470 nm, 48 lm) placed about 40 mm from the animal. Fluorescence was imaged using a long wavelength pass filter Schott GG495, (UQG Optics Limited) to block reflection of the excitation LED.

### Image capture and processing

A Canon Rebel T3i camera and Canon 100 mm macro lens mounted on a copy stand were used to obtain images of the entire animal. Higher magnifications, required to follow moving blood cells, was obtained using a Meiji stereoscopic microscope and the Canon camera. Images were processed using Adobe Photoshop software with any image enhancement applied uniformly to the entire frame.

## Results

The transparency of *C. inflata* makes it easy to see internal structure, but only if the structure is not itself transparent. Optimal observation is thus aided by a dye to create contrast. The dye neutral red is most commonly used in tissue culture as a stain specific for living cells, where it is transported into lysozymes to give the culture a red-pink color (Borenfreund and Puerner 1984). However, it also stains nuclear chromatin (Espelosin and Stockert 1982). Because it is non-toxic and bound complexes are stable it has been used to stain entire marine invertebrates for up to 7 days (Drolet and Barbeau 2006). In the present study neutral red was used as a relatively non-specific stain with stained blood cells identified by their motion. Moving blood cells define the vessels or channels that distribute blood, and the velocity of the cells reveals the dynamics of the circulatory system. Alternatively, injection of fluorescent dextran into the circulatory system while the heart is pumping identifies much of the vascular system that is imbedded in structures not involved in blood circulation.

In this report the terms “vessel” and “vascular” are used only to specify pathways or channels through which blood flows. There is no intent to suggest that a “vessel” has the histology of a blood vessel in a vertebrate, as the methods and equipment used in this study do not resolve structure at this scale.

### Structure of the tunicate and location of major blood vessels

The right side of a typical *C. inflata* stained with neutral red is seen in Fig. 1, panel A1, and diagrammed in panel A2. Tunicates have an inner cylindrical branchial basket attached along the ventral edge (bottom of panels) to the outer cylindrical body. Water is pumped through the intake, or oral, siphon at lower right, by cilia in the branchial basket. Food is trapped in a mucus net produced by the endostyle, pulled across the interior of the branchial basket and then drawn down into the stomach. Seawater escapes via the exit, or atrial siphon, just above the intake siphon. *C. inflata* has an enlarged exit siphon which in this image contains fecal matter, but is also used to contain and protect the juvenile tunicate “tadpoles”. The endostyle is the dark band along the ventral (lower in this panel) side of the animal where the branchial basket and the outer body join. The stomach, ovaries, and testis are the dark mass at the left end. The heart is the vertical band along the right edge of the visceral mass.

**Fig. 1.**
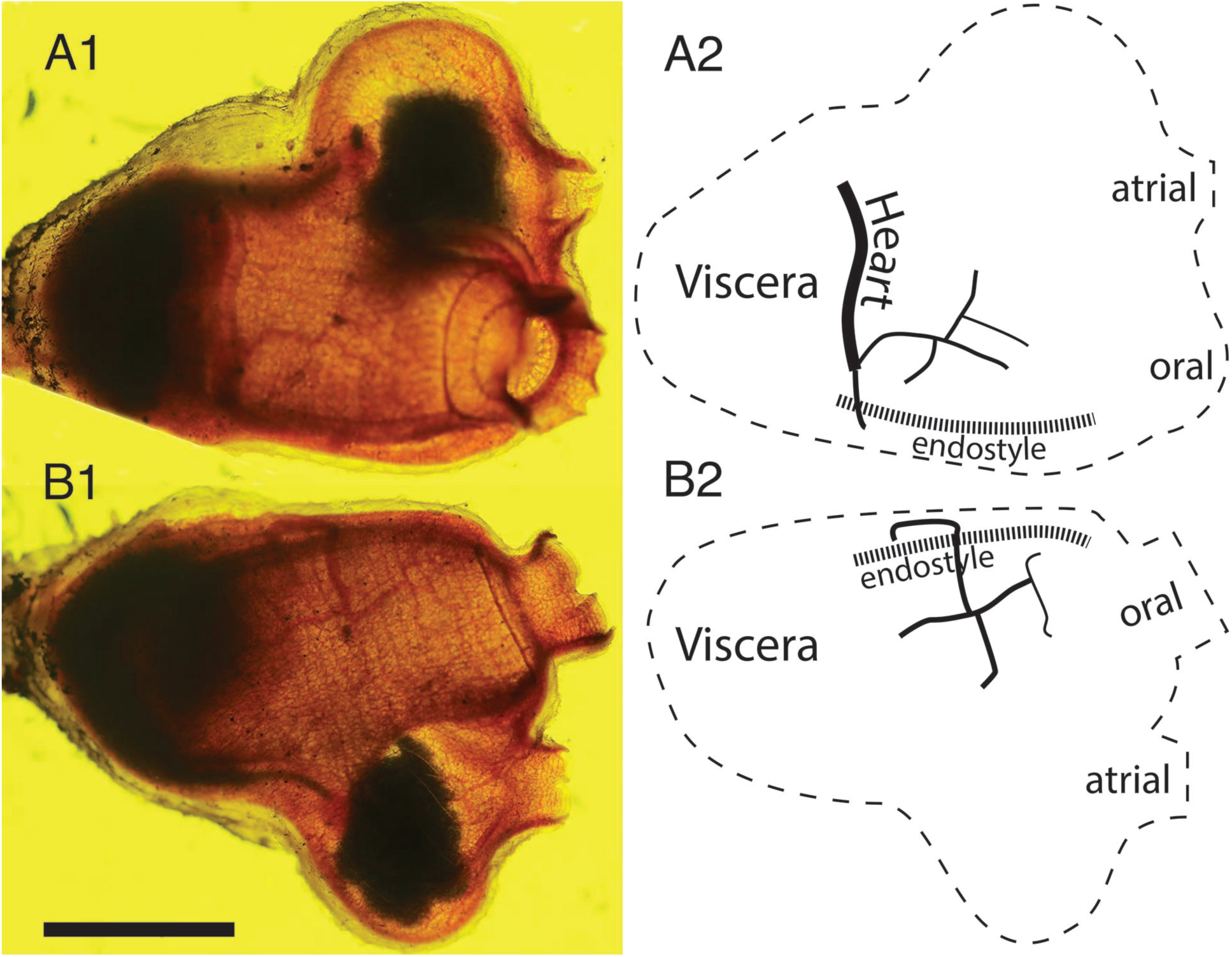
Blood vessels of a tunicate stained with neutral red. **A1.** Right side with ventral at bottom, anterior at the right: the heavily stained viscera (stomach, intestine, ovary, testis) fill the posterior, or left end. The heart is the translucent ribbon at the right side of the viscera and the endostyle is the red band along the ventral edge of the animal. **A2.** Diagram of tunicate in A1: oral (intake) and atrial (exit) siphons are at the right. The thick dashed line along the bottom of the tunicate is the endostyle. Medium and thin solid black lines indicate vessels leaving the ventral end of the heart. **B1.** Left side of tunicate in A1 rotated 180 degrees about the horizontal: endostyle now at top, viscera remain on the left, but hide the heart in this view. The vessel leaving the lower end of the heart in panel A1 is seen here at the top of the tunicate, looping over the endostyle and then branching to supply blood to the left side of the branchial basket. Scale bar at lower left represents 10 mm. **B2.** Diagram of tunicate in panel B1: the approximate volume of the animal is the product of the area of a triangle along its edges (270 mm^2^) and its thickness (10mm, data not shown) or 2700 mm^3^.

In *C. inflata* two large vessels exit the lower (ventral) end of the heart and pass to the middle of each side of the branchial basket where they repeatedly branch and merge into a rectangular net of vessels. One vessel connects to the middle of the right half of the basket (Fig. 1, panels A1, A2). The other passes over the endostyle to the left side of the animal and connects to the left side of the branchial basket (Fig. 1, panels B1, B2).

### Global pattern of blood flow

Moving blood cells define the functional vessels that are diagrammed in Fig. 2 and seen in Movie 1. The peristaltic heart of *C. inflata*, like other tunicates, reverses the direction of pumping every few minutes. During the abvisceral phase of heart action, seen in Movie 1, blood is drawn from the viscera into the posterior end of the heart to then flow out from the anterior end into the branchial basket via the two vessels mentioned previously. One vessel leaves the heart and then bends to the right to join the branchial basket on the right side of the animal. The vessel that carries blood to the left side of the branchial basket is not seen in the perspective of Fig 2. Blood that enters the middle of the right side of the branchial basket flows in a rectilinear network to vessels at both the dorsal and ventral edges of the basket, and then flows in a posterior direction toward the viscera to complete the circulation. A small fraction of blood leaving the heart supplies the mantle, but this vessel system is not apparent in the focal plane of Movie 1.

**Fig. 2.**
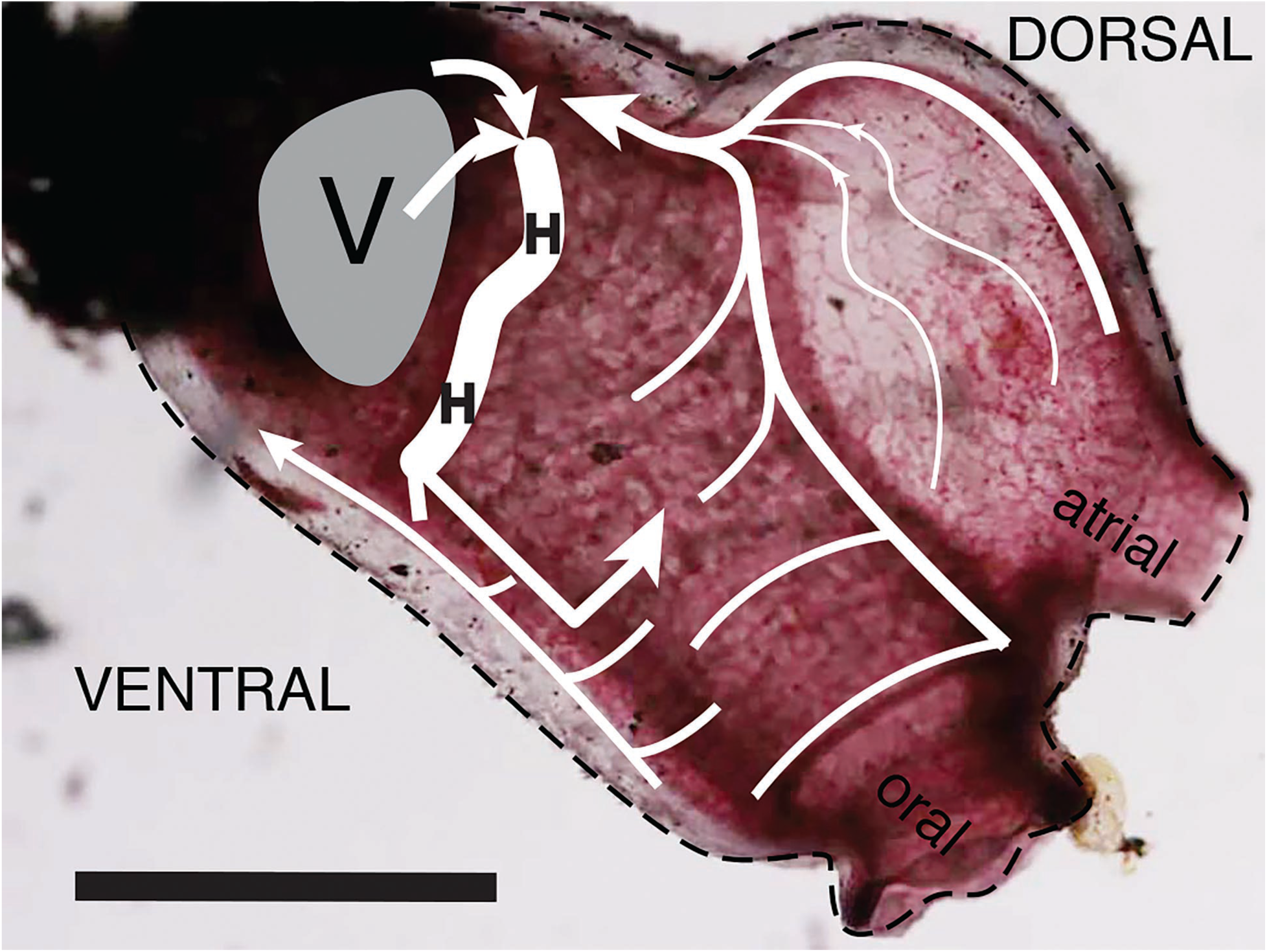
Blood flow in the right side of the tunicate. In this diagram the ventral side of the tunicate is down and the anterior end is to the right, as in Fig.1A. Blood leaves the heart at its ventral end and immediately splits into two vessels. Only the one supplying the right side of the branchial basket is shown here. After flowing through the branchial basket blood is collected by ventral and dorsal vessels and flows in the posterior direction into the viscera. Blood is collected from the viscera in a network of small vessels that progressively merge to form two large vessels that enter the heart at its dorsal end. When the heart pumps in the posterior direction all flows are reversed.

**Moive 1.** Blood flow in the right side of the tunicate oriented as in Fig. 2. The heart tube is the vertical stripe about one-third from the left side of image but to the right of the dark viscera. Peristaltic contractions moving downward force blood cells into vessels leading to the two sides of the branchial basket. There is always a constriction (clear segment) in the heart tube, and thus blood does not flow “backward” during a heartbeat.

Injection of the non-binding tracer, fluorescent dextran, into the circulation, allows direct visualization of the heart and the larger vessels. Dextran injected into the posterior end of the heart as blood is pumped toward the anterior filled a large vessel leaving the anterior end of the heart that connects to the right side of the branchial basket and spreads into a rectilinear vascular network in the basket wall (Fig. 3A). Dextran flowing to the left half of the branchial basket via the other vessel leaving the heart cannot be seen in this view. In another experiment injection of dextran into a vessel at the anterior end of the heart as blood is pumped in the posterior direction fills two large vessels that leave the heart from the posterior end (Fig. 3B). One vessel bends sharply in the ventral direction and then branches to supply the stomach and gonads. The other vessel branches toward the intestine and the main dorsal vessel. The initial injection also introduced fluorescent dextran into the pericardial sac that encompasses the inner heart tube. Thus the heart appears to be filled with fluorescent dextran even though at the time this image was obtained most of the fluorescent dextran in the interior heart tube has moved to the visceral region.

**Fig. 3.**
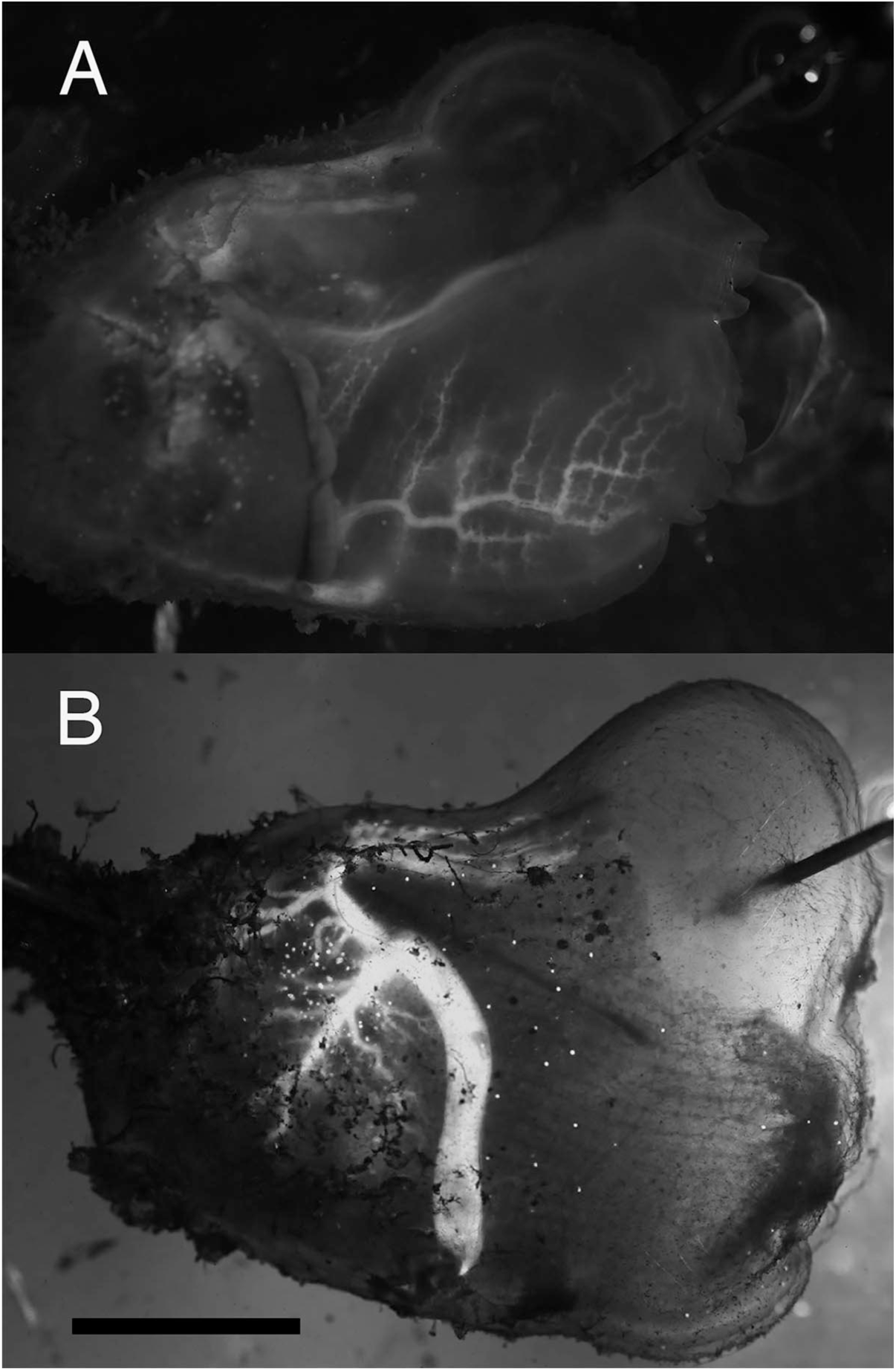
Florescent dextran infused into vessels. **A.** In the branchial basket. The tunicate is oriented as in Fig. 1. Dextran was injected (25 uL over 1.9 min) at the posterior end of heart while the heart pumped in the anterior (abvisceral) direction. This image was obtained 3.5 min after end of injection. Dextran inters the branchial basket through a branching tree of vessels in the side of the basket, not along an edge. **B.** In the viscera. Dextran was injected (15 uL over 28 sec.) at the anterior end of heart while heart pumped in the posterior (advisceral) direction. The image was obtained 6.5 min after end of injection. Dextran leaves the heart in two large vessels that then branch repeatedly to supply the visceral region, e.g. stomach and gonads. Some blood flows into the dorsal vessel and then moves toward the anterior. Scale bar at bottom left represents 10 mm.

### Heart structure and function

The *C. inflata* heart reverses direction of peristaltic contractions periodically as has been observed in other tunicates. Constrictions transverse the heart in an average of 1.4 seconds and approximately every 4 minutes, or 180 beats, the direction of movement reverses (Table 1).

**Table 1.** Six consecutive heart beat phases of the peristaltic heart. Advisceral flow is from the heart to the viscera while abvisceral flow is the reverse direction. The interior of the heart tube has a diameter of about 1.6 mm and length of 12 mm, to give a volume of 24 mm^3^. Since the average number of beats for one cycle is about 270, the average volume pumped in one cycle would be 6480 mm^3^ if the heart functioned as a piston moving down a tube with constant diameter. However, the space carrying the blood is more like an elongated oval between the two twists in the heart tube. Thus the actual volume pumped in one cycle is likely to be several-fold lower.

| **Direction** | **Time (sec)** | **Total beats** | **Beats / sec** |
| --- | --- | --- | --- |
| advisceral | 236 | 176 | 0.75 |
| abvisceral | 318 | 220 | 0.69 |
| advisceral | 243 | 160 | 0.66 |
| abvisceral | 293 | 197 | 0.67 |
| advisceral | 261 | 183 | 0.70 |
| abvisceral | 252 | 170 | 0.67 |
| **ad: average** | **247** | **173** | **0.70** |
| **ab: average** | **288** | **196** | **0.68** |

The heart consists of two concentric cylinders fused at the ends. The inner cylindrical tube contracts and twists progressively along its length to produce a moving constriction that forces the blood from one end to the other. The surface of the outer tube is relatively static, and thus it does not directly participate in the pumping process. The pericardial space between the tubes is filled with fluid. Fluorescent dextran injected into the pericardial space of the *C. inflata* heart clearly defines the tube enclosing the heart (Fig. 4). Dextran remained in the pericardial space during a 24-hour period when the heart was beating at its normal rate. Thus, this space is quite isolated from the rest of the animal.

**Fig. 4.**
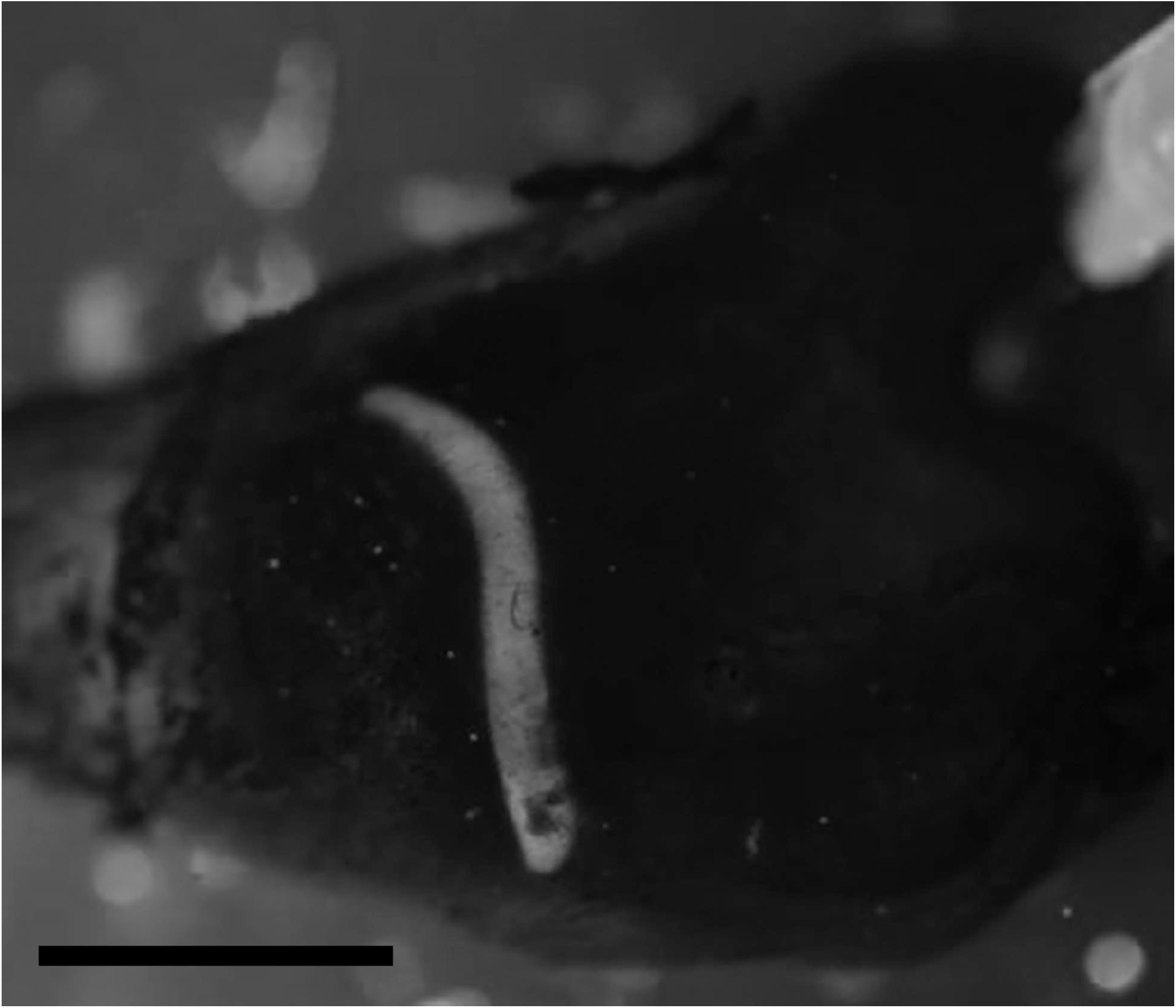
Fluorescent dextran infused into the pericardial space. This image was obtained more than 24 hours after the injection and the heartbeat was normal throughout this period. There is no evidence of a constriction in the middle of the pericardial tube. As in previous figures ventral is at bottom and anterior is to the right and the scale bar at bottom represents 10 mm.

The flow of stained blood cells enables contractions of the heart tube to be followed in some detail. As one peristaltic constriction approaches the end of the heart, another constriction forms at the other end and thus the heart tube never appears open. The constriction that moves the blood through the heart tube does not seem to be cylindrically symmetric, but rather appears as a helical fold that moves along the tube. About halfway along the heart tube is a stationary pinch of about half the average diameter of the heart tube. As the fold approaches this pinch, blood is forced through and a new fold appears on the other side with mirror image twist. Thus the pinch does not appear to function as a valve but rather as a support for the inner heart tube. The anterior (ventral) half of the heart, the heart-tube folds, and the vessel from the heart to the right side of the branchial basket is seen in Fig. 5A, and diagrammed in Fig. 5B. Movie 2, which has the same field of view as Fig. 5, documents the movement of the heart folds.

**Fig. 5.**
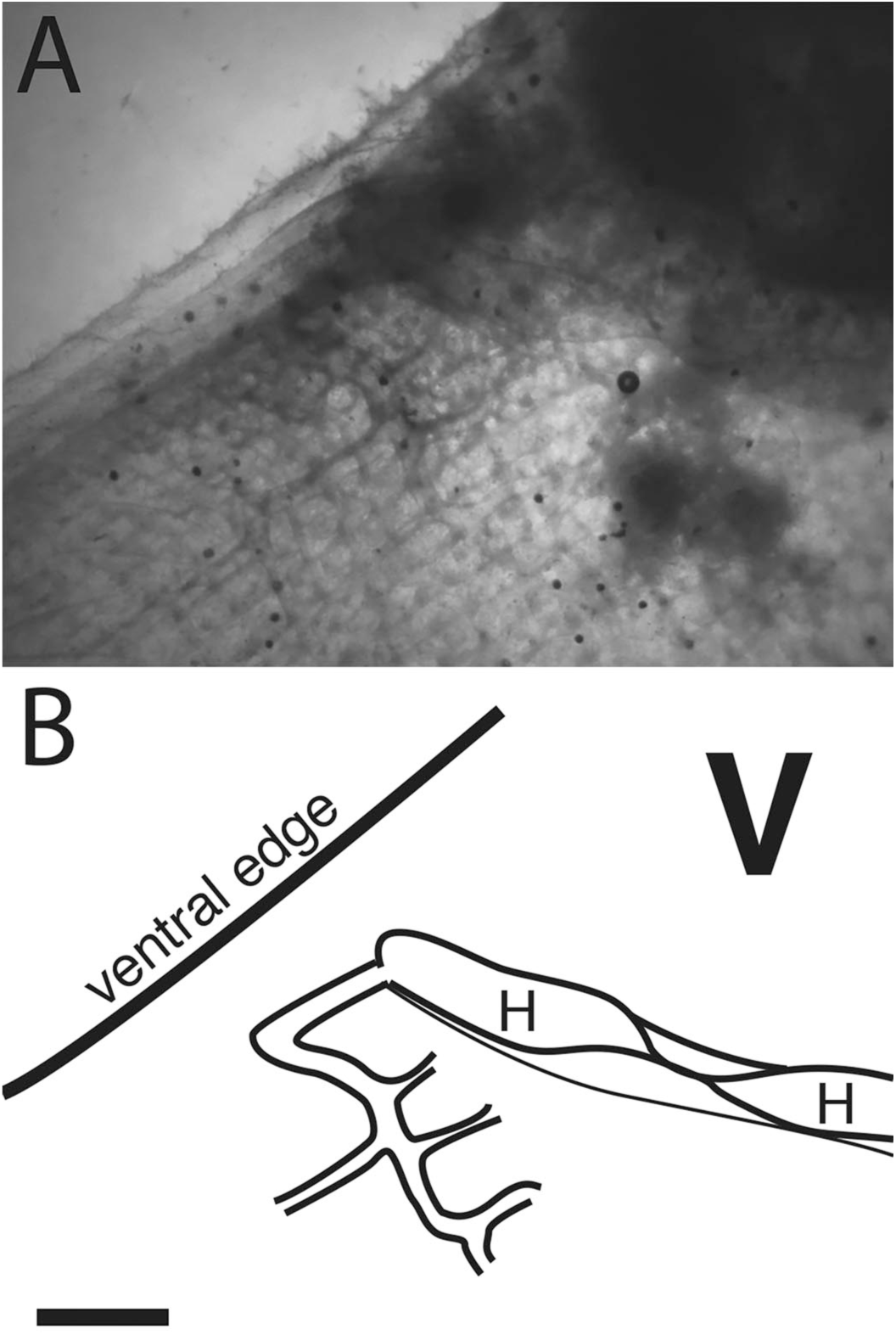
Folds in the internal heart tube at the central constriction. At the top is a diagram of the ventral end of the heart and the vessel that connects to the right side of the branchial basket. The tunicate is rotated relative to previous figures, and the ventral edge is upward, anterior to the left. The “V” at upper right indicates the viscera. Segments of the heart tube containing blood on each side of the twist are labeled “H”, and the scale bar represents 2 mm. The bottom panel is a photo of this region in a tunicate stained with neutral red.

**Moive 2.** Folds in the internal heart tube at the central constriction. The orientation and scale of this movie is described in Fig. 5.

Rapidly moving stained blood cells can be seen in the heart, but the high velocity generates streaks in video frames, and cells come in and out of focus due to the complex flow pattern in the heart. However, it was possible to follow the position of individual blood cells in a large vessel just anterior to the heart as seen in Fig. 5, and thus to determine the velocity versus time profile (Fig. 6). The velocity curve is approximately sinusoidal with a period slightly more than a second, consistent with Table 1. There is a reproducible small dip in the region of maximum velocity. The velocity is always positive, i.e. cells always move in the direction of the heartbeat, but the minimum velocity is only about a tenth the maximum value. In a typical animal (Fig. 1), the heart is about 12 mm long, and a constriction moves through the heart every 1.4 seconds, giving an average velocity of 9 mm s^-1^ for the constriction. The peak velocity of about 2 mm s^−1^ in the vessel just anterior to the heart (Fig. 6) is thus several-fold lower than the velocity of the heart constriction. However, this is but one of several large vessels the heart empties into, and the total cross sectional area of these vessels can only be estimated.

**Figure 6.**
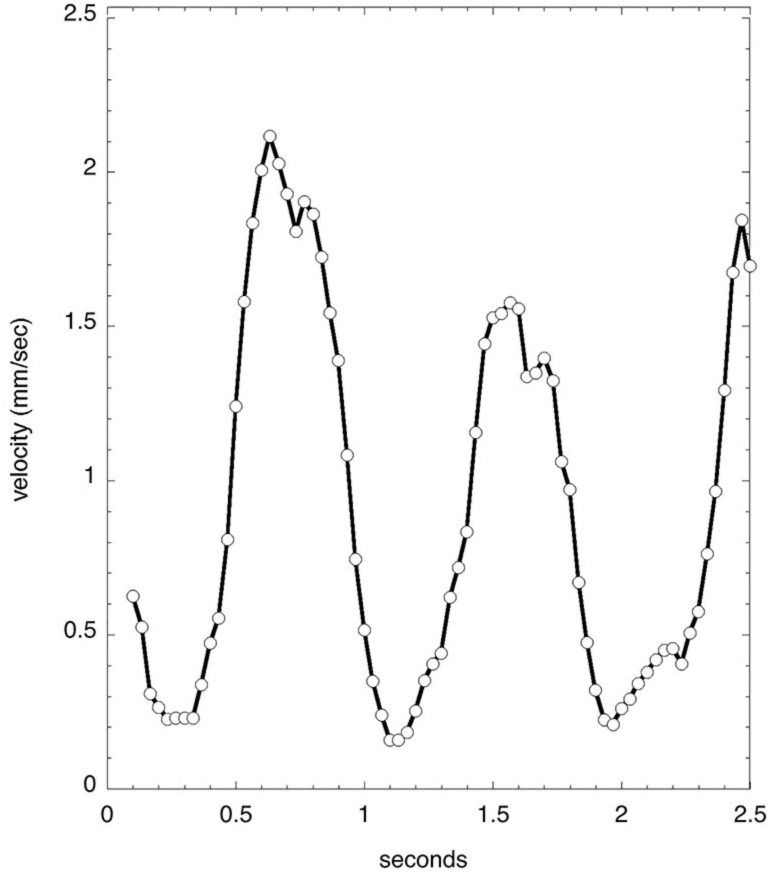
Velocity of a blood cell in a large vessel. A movie with approximately the same field of view as Fig. 5 was disassembled into individual frames (30 for each second), and the position of one densely stained cell was followed through sequential frames. Velocity values in the graph are moving averages using values from six consecutive frames. The average blood cell velocity over two complete heartbeats (0.20 to 1.97 sec) was 0.88 mm/sec.

Blood cell velocities in three small vessels in the mantle at the extreme anterior end of the animal are much smaller that seen in the large vessel (Fig 7). These velocities appear stochastic, with an average of about 0.3 mm s^−1^, but all are positive (move in the same direction) throughout several heartbeats.

**Figure 7.**
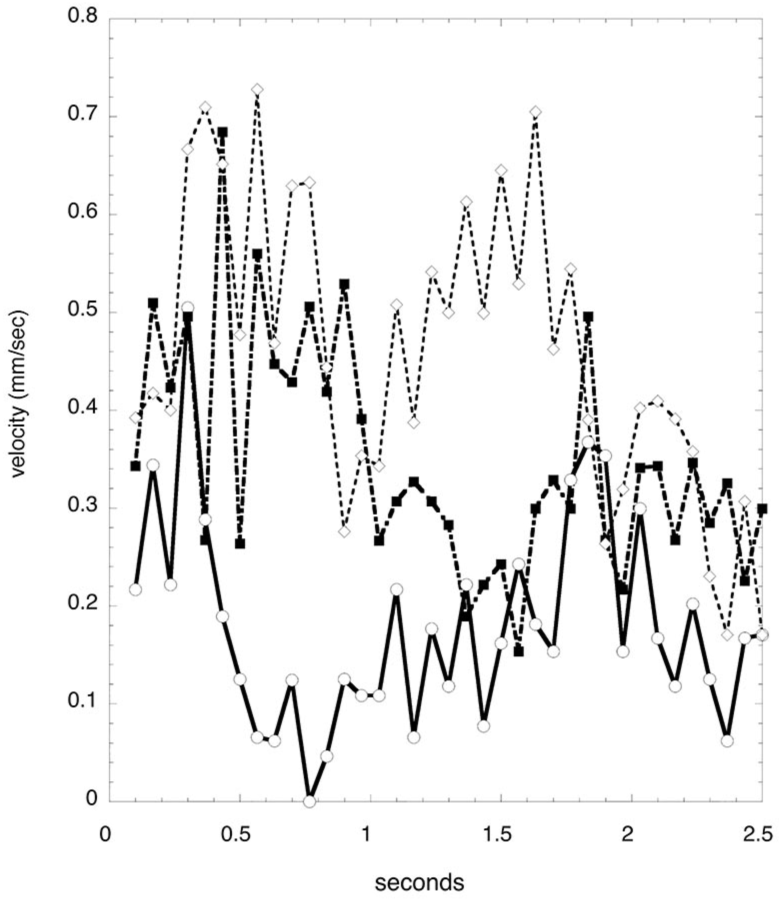
Velocity of blood cells in three small vessels. A movie of stained blood cells in the mantle surrounding the exit siphon was analyzed in the same manner as described in Fig. 6.

### Blood flow in the branchial basket

The cylindrical branchial basket occupies a major portion of the interior volume of the tunicate. As described in a previous section and seen in Fig. 3A, blood flows from the ventral end of the heart to the middle of each side of the branchial basket. On the interior of the basket a rectangular network provides support and distributes blood (Fig. 8, panel A). Blood then flows into an exterior layer perforated with pairs of spiral stigmata resembling the Ionic volutes of Greek columns (Fig. 8, panel B). Cilia line the inside edges of the stigmata, but can only be seen easily by their motion. Fluorescent dextran injected into the blood circulation reveals the structure of the duct containing the stigmata in low resolution (Fig. 8, panel C). Blood cells can be seen moving in the area immediately adjacent to and between the stigmata. This is consistent with stigmata being openings through a double layer of cells, and blood circulating in the space between the cell layers.

**Figure 8.**
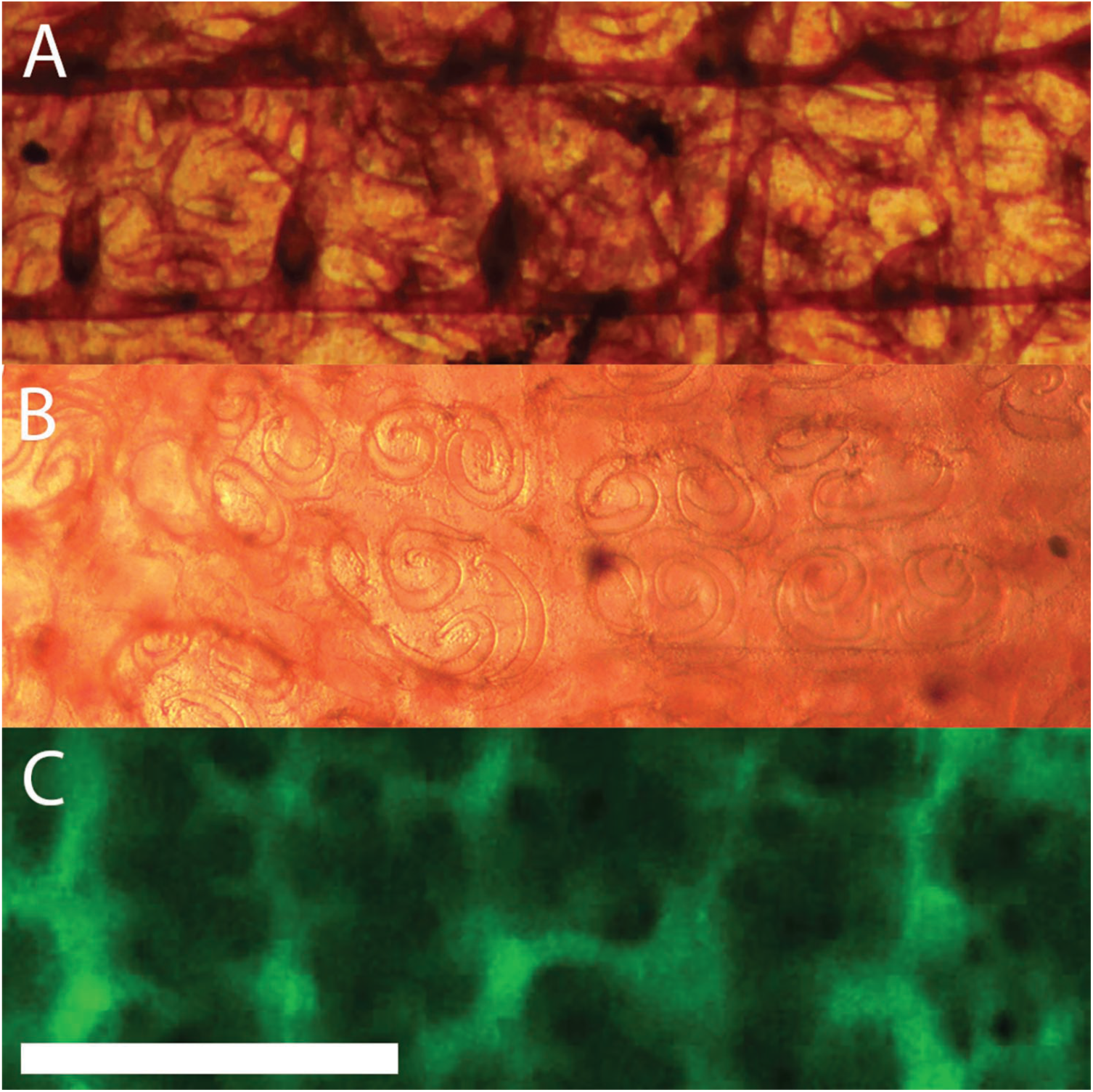
Three layers of structure in the branchial basket. Axis of the tunicate is horizontal with anterior to the right (same orientation as in other figures of this publication). Height of view in each panel is 1 mm. Each panel is from a different animal, but with corresponding position and orientation. **A.** Interior surface (stained with neutral red). The two prominent horizontal lines in the figure (vertical in the animal) are the longitudinal bars. There are 5 vertical (lateral in the animal) branchial vessels associated with transverse bars that lie under the longitudinal bars (only the two on the right are clearly defined in this image). **B.** Exterior surface (stained with neutral red). The spiral worm-like shapes that resemble Ionic volutes are the openings (stigmata) through which water is pumped by cilia along the edges of the openings. Movies at a higher magnification reveal the motion of beating cilia on the edge of the stigmata. **C.** Exterior surface after injection of fluorescent dextran posterior to heart. Blood flows through a complex network of ducts linked together with net circulation perpendicular to the axis of the tunicate. Scale bar at bottom left represents 1 mm.

Blood from the branchial basket is collected by vessels along the ventral and dorsal edges of the branchial basket to flow to the posterior of the animal (summarized in the schematic of Fig. 2). When the heart reverses direction the direction of blood flow in all parts of the animal reverses.

## Discussion

The movement of stained blood cells and the pattern of fluorescent dextran injected into blood vessels have defined the heart, a network of vessels in the body, and ducts in the branchial basket. As in other tunicates the heart is peristaltic and periodically reverses direction (Newman 1939). Durations of flow in the two directions are similar, however the abvisceral (viscera to heart to branchial basket) direction is seen in vertebrates, e.g. fish, and is used by Millar (Millar 1953) in describing blood flow in the tunicate *Ciona.* Thus the abvisceral direction will be assumed in this discussion unless otherwise specified.

### Blood flow

Dextran distribution and cell flow are complimentary tools, dextran clearly defining the larger vessels while blood cell flow reveals medium and small vessels and allows the determination of blood velocity. The circulation of blood can be followed through most of the tunicate body. Blood leaves the anterior end of the heart in two vessels; one connects to the right side of the branchial basket and the other crosses over the endostyle to connect to the left side of the branchial basket. The blood is then distributed through both the longitudinal and transverse bars (or tubes), into ducts containing the stigmata. Blood is collected by one large vessel on the ventral and another on dorsal side of the body and then flows into the visceral region. Blood is collected from the visceral region by a branched vascular tree that becomes two large vessels entering the posterior end of the heart. It has not been possible to trace blood flow through the visceral region, specifically in the space between the posterior ends of the dorsal and ventral vessels and the ends of the vascular tree that feeds blood into the heart. A major difficulty in following circulation is the density and opacity of the organs in this region. It is also possible that blood does not always flow through clearly defined vessels, but rather through a network of cavities as seen in the liver and spleen of vertebrates (Richardson and Granger 1984).

### Vessel structure

At the modest magnifications used in this study it was not possible to see the structure of vessels at the cellular level. However, the two large vessels that leave the anterior end of the heart to follow asymmetric paths to the two sides of the branchial basket do not appear to be associated with any of the symmetric structures in that region. Thus in that sense they are independent structures.

The molecular integrity of vessels can be inferred by the rate infused fluorescent dextran leaks into surrounding tissue. A halo of fluorescence can be seen around the large vessels supplying blood to the branchial basket about 5 minutes after the start of the injection (Fig. 3, panel A). However, since these vessels supply blood to smaller vessels and ducts, much of the halo seen at this resolution could represent distribution, not leakage. Dextran flowing into the posterior region (Fig. 3, panel P) remains in the major vessels for more than 6 minutes after the injection. Dextran of this molecular size has been used to follow plasma leakage in comparable times from vessels in the hamster cheek pouch induced by leukotrienes (Dahlen et al. 1981). However, in these studies leakage could be compared in normal and treated animals while in this report I can only make absolute statements about vessel permeability. Perhaps all that can be concluded is that tunicate vessels can contain dextran for at least five minutes, a time comparable to vertebrate vessels. However, blood pressure in tunicates is about 100 times lower than that in vertebrates (Jones 1985), thus absolute tunicate vessel permeability could be larger and give comparable leakage rates.

### Heart structure and function

The slightly curved peristaltic heart extends across most of the left side of the body at the anterior edge of the visceral region and is hidden by the visceral mass when viewed from the right side. The heart is joined to two large vessels at both posterior and anterior ends. A peristaltic heart tube branching into two vessels at each end, is the canonical structure of the vertebrate embryonic heart soon after the heart tube is formed (Gilbert 2010; Santhanakrishnan and Miller 2011).

The diameter of the outer pericardial tube is approximately constant over its length, but the inner heart tube has a fixed constriction approximately in the middle. A change in the twist of the heart tube as the peristaltic contraction passes through this constriction implies it is an anatomical feature with possible function, however it is not a valve. Hecht reported seeing a constriction, or node, in the middle of the heart of *Ascidia atra* (Hecht 1918), a tunicate in the same order as *C. inflata.* He described the node in detail, and stated it was a “real landmark in the structure”. However, Goodbody, observing the heart of *Corella willimeriana,* also observed an apparent constriction in the heart but believed it was not a real anatomical structure but just a “shifting of the position the ralph”, (Goodbody 1974). He referred to Hecht’s report and stated that Hecht also thought the node was just a shifting of the ralph. In my opinion this is not an accurate description of Hecht’s claims.

Microscope optics, illumination, and animal species used by Hecht, Goodbody, and the present author are all different, and the two previous authors did not stain the animals. However, I have seen “the node” in all my observations, and along with Hecht believe it is a “real landmark”.

The present report does not propose a model for heart action but the existence of the middle node has potential meaning for the phylogenetic relation between tunicates and vertebrates. In the embryonic development of vertebrates a pinch in the middle of the heart tube is the beginning of differentiation into ventricle and atrial chambers, which have no obvious correlates in tunicates that periodically reverse direction of heart pumping. However, the pinch in the tunicate heart might be the relic of an ancestor with unidirectional blood flow.

The twist in the inner heart tube, which constrains the blood, moves across the heart with an average velocity of approximately 9 mm s^−1^. However, the blood moves through the heart as an elongated ellipsoid, or spindle, and thus the volumetric output is less than the product of the cross sectional area and the velocity. In one of the two large vessel near the anterior end of the heart blood cells have a semi-sinusoidal velocity-time curve with a maximum of 2.2 mm s^−1^ and a slight dip at the maximum. It is possible that this dip is due to the constriction in the middle of the heart tube. It is difficult to measure the diameters of the heart and vessels with much accuracy since the edges do not appear sharp in the microscope. Because the area of a tube is proportional to the square of the diameter, errors in the calculated area are twice those of the diameter measurements. Cells move in one direction throughout the beat, consistent with the presence of at least one twist in the heart tube throughout the heartbeat so blood does not leak in the reverse direction. For vessels in the mantle maximum velocities range from 0.7 to 0.5 mm s^−1^, but velocities are irregular and not sinusoidal.

The fish is perhaps the best vertebrate comparison, and a blood velocity of 1.7 mm s^−1^ has been measured in the aorta of a 5 day-post fertilization Zebrafish (Watkins et al. 2012). Zebrafish of this age are only about 2.5 mm long (Parichy et al. 2009) compared to a typical length of 30 mm for the Corella in this study, and blood velocity increases as the fish grows. However, blood velocities in the tunicate and fish are at least roughly comparable.

### Blood flow in the branchial basket

The branchial basket is the largest and most conspicuous structure in most tunicates and its structure and pattern of vessels are the basis of division of the tunicate class *Ascidiacea* into orders. The wall of the branchial basket consists of two parallel layers of cells, penetrated by openings, the stigmata, through which water is pumped by seven rows of cilia bearing cells along the circumference of the stigmata (Burighel and Cloney 1997). The wall of the branchial basket is thus similar to the double-layered lamella (Olson 2002), that constitutes the gills of fish. However, in the fish the layers are closer together, are held in place by pillar cells, and of course are not penetrated by stigmata.

Blood, supplied by transverse and longitudinal bars (tubes), seen in Fig 8A, flows between the walls, Fig 8C, and is collected by vessels along the ventral and dorsal edges of the basket. Thus, the shapes of the stigmata, which are spiral in *C. inflata*, define not only the flow of water through the wall, but also the flow of blood within the walls of the branchial basket. In *C. inflata* the two vessels from the heart to the middle of the sides of the basket, the transverse and longitudinal bars, and finally the hollow walls of the branchial basket and the stigmata should all be considered part of the pattern of blood circulation characteristic of the species.

### Comparison with the circulatory system of *Ciona intestinalis*

The most studied tunicate is *C. intestinalis* and the most comprehensive general description of the anatomy and histology of this tunicate is arguably the 124 page monograph by Millar (Millar 1953) that combines microscopic observation of live animals, stained sections, and sections from animals in which the vascular system have been filled by dyed latex.

In C. *intestinalis* there appears to be no constriction in the middle of the heart tube, however it would be difficult to see one since the heart bends sharply in the middle more than 90 degrees. Blood flows from the anterior end of the heart directly into the vessel (or channel) running along the ventral edge of the endostyle. Then parallel transverse bars transport blood to the basket that drains into the dorsal vessel and down to the posterior of the tunicate. Thus, there are no blood vessels independent of the branchial basket in this part of the circulatory system. The stigmata of *C. intestinalis* are thin rectangles aligned with the longitudinal bars. Millar did not describe the double walls of the branchial basket, and thus may have been unaware of this feature.

Millar states: “The blood vessels of *Ciona* are channels and lacunae in the connective tissue and are not lined with endothelium, except close to the ends of the heart…” However, he does not state which vessels he has examined, or the criteria used for identification of endothelia.

In Text-Fig. 6, pg 57 Millar shows blood flowing from the heart through the ventral endostyle, across the branchial basket, and back down the dorsal vessel to the visceral region; a sequence that suggests a circulation. He sites Von Skramlik (Skramlik 1929) as finding that there is “true circulation of the blood”, and does not state opposition to this conclusion.

### Comparison with circulation in a colonial tunicate

The circulatory system of the colonial tunicate *Botryllus schlosseri* has been studied in some detail (Burighel and Brunetti 1971). Each zooid in the colony has an internal structure similar to that seen in *C. inflata* and *C. intestinalis*. The blood appears to circulate in a path with the same topology described above for solitary tunicates, although there is at least one small vessel at the end of each zooid that connects to a colony wide vessel net.

### Open or closed, tidal or circulating

Discussion of blood circulation in tunicates is complicated by potential confusion between two distinct characteristics of a circulatory system and by the use of terms that are not neutral with respect to these characteristics.

In open circulation blood does not flow through vessels that are distinct and isolated from the cells of the body tissues, rather the tissues are directly bathed in the blood, which is then called hemolymph (*hemo,* blood; *lymph,* fluid cells are bathed in). In a closed system blood is confined to a distinct vascular space. Thus the use of “blood” in the title of this report suggests circulation in *C. inflata* is closed and confined to a vascular system. However, while blood in a closed circulation is confined, water, blood proteins and white blood cells percolate at various rates through vessel walls to become lymph. Thus, open versus closed may best be considered a quantitative parameter describing the degree or rate blood is filtered before reaching the cells. In this report the persistence of dextran in the vasculature of *C. inflata* is comparable to that seen in the closed circulation of vertebrates, which suggests that circulation in this tunicate can also be described as closed. This is not the majority opinion for tunicates since a recent review states that “tunicates are generally considered to have an open circulatory system” (Davidson 2007).

The endothelial cells that line or constitute the blood vessels of vertebrates are considered the essential barrier between blood and lymph (Reiber and McGaw 2009). Since most invertebrates do not have endothelial cells, and Millar reports that endothelial cells are found only near the heart in the tunicate *C. intestinalis,* one might conclude that circulation in tunicates must be open. However, Reiber and McGaw argue that the open-closed character of circulation is a continuum, note that some invertebrates that lack endothelial cells have a functional closed circulation, and that the invertebrate cephalopod molluscs have a vascular system lined with endothelial cells (Reiber and McGaw 2009).

Circulation can be used as a generic term for the flow of blood or hemolymph. However, used more specifically circulation is a flow that moves around a closed circuit. Blood circulates in through a network of circuits in vertebrates. The hearts of tunicates, including *C. inflata,* pump in one direction for a period and then reverse for a similar time. An early text on tunicates states: “blood does not go around a true circuit but shuttles back and forth through the heart” (Newman 1939). This quote describes a tidal or reciprocating flow, where blood is pumped from one compartment to another and then back through the same path.

However, observation of blood cells in *C. inflata* reveal they move in circuits, as seen in Movie 1, although the flow in all the circuits change direction when the heart changes its pumping direction. The apparent circulation-tidal contradiction is resolved in the case of *C. inflata* by consideration of the volume of blood pumped by the heart during one directional phase, which can be done either using the volume of the peristaltic heart and the number of beats per directional phase, or integrating the velocity of blood cells in a large vessel near the heart over a directional phase. The total volume over the 4 minutes the heart pumps in one direction is larger that the total volume of the animal (tissue and enclosed water), and thus much larger than the total volume of blood. Thus, blood must circulate in this tunicate in the strict sense, but in opposite directions during the two directional phases of heart action. A similar calculation and conclusion has been made by Kriebel for circulation in the tunicate *Ciona intestinalis* (Kriebel 1968).

If reversal of blood flow is not required by the anatomy of the circulatory system, there may be a physiological advantage in periodic reversal. One possibility is that the gradient of oxygen and nutrients along the circulation pathway is sufficiently steep that tissues in the second half of the circulation loop do not receive optimal levels. Periodic reversal would provide a more uniform delivery, and Ruppert et al., on page 946 suggest this possibility (Ruppert et al. 2004). In addition, the peristaltic heart could alter the ratio of beats in one versus the other direction, and thus compensate for changes in nutrient or oxygen supplies and demands. However, no such regulation has been documented.

### Evolutionary relationships

It might be appropriate to compare tunicate blood circulation to circulation in the vertebrate fish, with emphasis on the differences in blood flow to the branchial basket in *C. intestinalis* and *C. inflata*. In the fish the end of the heart ventricle is sufficiently close to the median plane and distant from the gills that blood flows easily into a ventral vessel that branches symmetrically right and left to become the ventral ends of the gill arches (Olson 2002). This geometry is similar to that seen in *C. intestinalis*, where blood flows into a vessel under the endostyle and then symmetrically up left and right branchial arches.

The topology of circulation in *C. inflata* is the exception, both in that blood flows from the heart into the middle of the branchial basket, and that the vessel that supplies the left side of the branchial basket crosses over the endostyle. Knowledge of embryological development of the circulatory system in this region may explain the origin of the difference in topology.

The pattern and structure of stigmata is one feature used to define tunicate species, and van Name (Van Name 1945) present diagrams of stigmata for several *Corella* species (but not *inflata*) that are similar to the paired spirals described in this publication. In contrast, the stigmata of *Ciona intestinalis* are essentially simple rows of rounded rectangles. A correlation between the geometry of the vascular connection between heart and branchial basket and the structure of the branchial stigmata is an intriguing possibility.

DNA sequences of 18S rDNA genes suggest that the genera *Ciona* and *Corella* are as closely related as the two extremely similar *Ciona* species *intestinalis* and *savignyi* (Stach and Turbeville 2002). However, the complete genome sequences for *C. intestinalis* and *C. savignyi* suggested that the these two tunicates have diverged from a common ancestor at a time approximately equal to the divergence of the chicken and human (Berna et al. 2009). Thus available genomic sequence data provides little guidance in predicting differences in the topology of blood circulation in *Ciona* and *Corella*.

### Future studies

A comprehensive histological study of the blood vessels *C. inflata* and *C. intestinalis* would extend our understanding of the relation of the circulatory systems of tunicates to that of vertebrates. Studies of the blood circulation pattern in other tunicates are needed to determine how common a bilateral connection between the heart and the branchial basket is, and whether it is correlated with sigmoidal stigmata. The topology of blood flow into the branchial basket may represent an apomorphy that defines two clades of tunicates, or it may merely constitute one member in a spectrum of patterns.

Determination of the complete genome sequence of *C. inflata* would enable a comparison with the known genome sequence of *C. intestinalis*, with the possibility of identifying genes that control development of the circulatory system.

